# The plastid genome in Cladophorales green algae is encoded by hairpin plasmids

**DOI:** 10.1101/145037

**Authors:** Andrea Del Cortona, Frederik Leliaert, Kenny A. Bogaert, Monique Turmel, Christian Boedeker, Jan Janouškovec, Juan M. Lopez-Bautista, Heroen Verbruggen, Klaas Vandepoele, Olivier De Clerck

**Affiliations:** Department of Biology, Phycology Research Group, Ghent University, 9000 Ghent, Belgium; Department of Plant Biotechnology and Bioinformatics, Ghent University, 9052 Ghent, Belgium; VIB Center for Plant Systems Biology, 9052 Ghent, Belgium; Bioinformatics Institute Ghent, Ghent University, 9052 Ghent, Belgium; Botanic Garden Meise, 1860 Meise, Belgium; Institut de biologie intégrative et des systèmes, Département de biochimie, de microbiologie et de bio-informatique, Université Laval, Québec (QC) Canada; School of Biological Sciences, Victoria University of Wellington, Wellington, New Zealand; Department of Genetics, Evolution and Environment, University College London, London WC1E 6BT, United Kingdom; Department of Biological Sciences, The University of Alabama, Tuscaloosa, AL35484-0345, USA; School of BioSciences, University of Melbourne, Victoria 3010, Australia

## Abstract

Virtually all plastid (chloroplast) genomes are circular double-stranded DNA molecules, typically between 100-200 kb in size and encoding circa 80-250 genes. Exceptions to this universal plastid genome architecture are very few and include the dinoflagellates where genes are located on DNA minicircles. Here we report on the highly deviant chloroplast genome of Cladophorales green algae, which is entirely fragmented into hairpin plasmids. Short and long read high-throughput sequencing of DNA and RNA demonstrated that the chloroplast genes of *Boodlea composita* are encoded on 1-7 kb DNA contigs with an exceptionally high GC-content, each containing a long inverted repeat with one or two protein-coding genes and conserved non-coding regions putatively involved in replication and/or expression. We propose that these contigs correspond to linear single-stranded DNA molecules that fold onto themselves to form hairpin plasmids. The *Boodlea* chloroplast genes are highly divergent from their corresponding orthologs. The origin of this highly deviant chloroplast genome likely occurred before the emergence of the Cladophorales, and coincided with an elevated transfer of chloroplast genes to the nucleus. A chloroplast genome that is composed only of linear DNA molecules is unprecedented among eukaryotes and highlights unexpected variation in the plastid genome architecture.

## Introduction

Photosynthetic eukaryotes originated more than 1.4 billion years ago following an endosymbiotic event in which a heterotrophic ancestor of the Archaeplastida engulfed a cyanobacterium that became stably integrated and evolved into a membrane-bound organelle, the plastid^1, 2^. Following this primary endosymbiosis, an intricate history of plastid acquisition via eukaryote-eukaryote endosymbioses resulted in the spread of plastids to distantly related eukaryotic lineages^3, 4, 5^.

Plastids have retained a reduced version of the genome inherited from their cyanobacterial ancestor. A core set of genes involved in the light reactions of photosynthesis, ATP generation, and functions related to transcription and translation is typically retained ^6, 7^. Many genes have been lost or transferred to the nuclear genome and, as a result, plastids are dependent on nuclear-encoded, plastid-targeted proteins for the maintenance of essential biochemical pathways and other functions such as genome replication, gene expression, and DNA repair^8^. Despite the wide diversity in size, gene content, density and organisation of plastid genomes among different eukaryotic lineages^9, 10, 11^, nearly all plastid genomes consist of a single circular-mapping chromosome, typically between 100-200 kb and encoding circa 80-250 genes^7, 12^.

While fragmented mitochondrial genomes evolved several times independently during the evolution of eukaryotes^11, 13^, fragmented plastid genomes are only known in dinoflagellates^14^ and a single green algal species^15^. In peridinin-containing dinoflagellates, the chloroplast genome is fragmented into plasmid-like DNA minicircles of 2-3 kb, most of which carry one gene only^14, 16^. Larger minicircles of up to 12 kb have also been described^17^, as well as minicircles containing two genes^18^, and ‘empty’ minicircles without genes^19, 20^. The genes located on these minicircles mostly encode key components of the major photosynthetic complexes, including subunits of photosystems I and II, the cytochrome b6f complex, and ATP synthase, as well as rRNAs and a few tRNAs^14^. The only other alga with a fragmented chloroplast genome is the green alga *Koshicola spirodelophila*, but here the level of fragmentation is minor: the plastid genome is divided into three large circular chromosomes totalling 385 kb, with a gene content comparable to other green algae^15^. In addition, plastid minicircles that coexist with a conventional plastid genome have been observed in a few algae, including dinoflagellates with haptophyte-derived plastids^21^ and the green alga *Acetabularia*^22, 23^.

Although plastid genomes generally assemble as circular-mapping DNAs, they can take multiple complex conformations *in vivo*, including multigenomic, linear-branched structures with discrete termini^24, 25^. The alveolate *Chromera velia* is the only known alga with a linear-mapping plastid genome with telomeric arrangement^26^, and is also atypical in that several core photosynthesis genes are fragmented. Linear plastid genomes, however, may be more widespread as several plastid genomes currently do not map as a circle^27^.

Currently, and in stark contrast to other green algae^9, 28, 29, 30, 31^, little is known about the gene content and structure of the chloroplast genome in the Cladophorales (Ulvophyceae), an ecologically important group of marine and freshwater green algae, which includes several hundreds of species. These macroscopic multicellular algae have giant, multinucleate cells containing numerous chloroplasts (Fig. 1a-c). Most attempts to amplify common chloroplast genes have failed^32^, with only one highly divergent *rbcL* sequence published thus far for *Chaetomorpha valida*^33^. An atypical plastid genome in the Cladophorales is suggested by the presence of abundant plasmids that have been observed in the chloroplasts of several species^34, 35^. These plasmids represent a Low Molecular Weight (LMW) DNA fraction, visible on agarose gels of total DNA extracts (Fig. 1d). Pioneering work revealed that these plasmids are single-stranded DNA (ssDNA) molecules of 1.5-3.0 kb that fold in a hairpin configuration and lack sequence similarity to the nuclear DNA^36, 37^. Some of the hairpin plasmids contain putatively transcribed sequences with similarity to chloroplast genes encoding subunits of Photosystems I and II (*psaB*, *psbB*, *psbC* and *psbF*)^36^.

**Figure 1.**
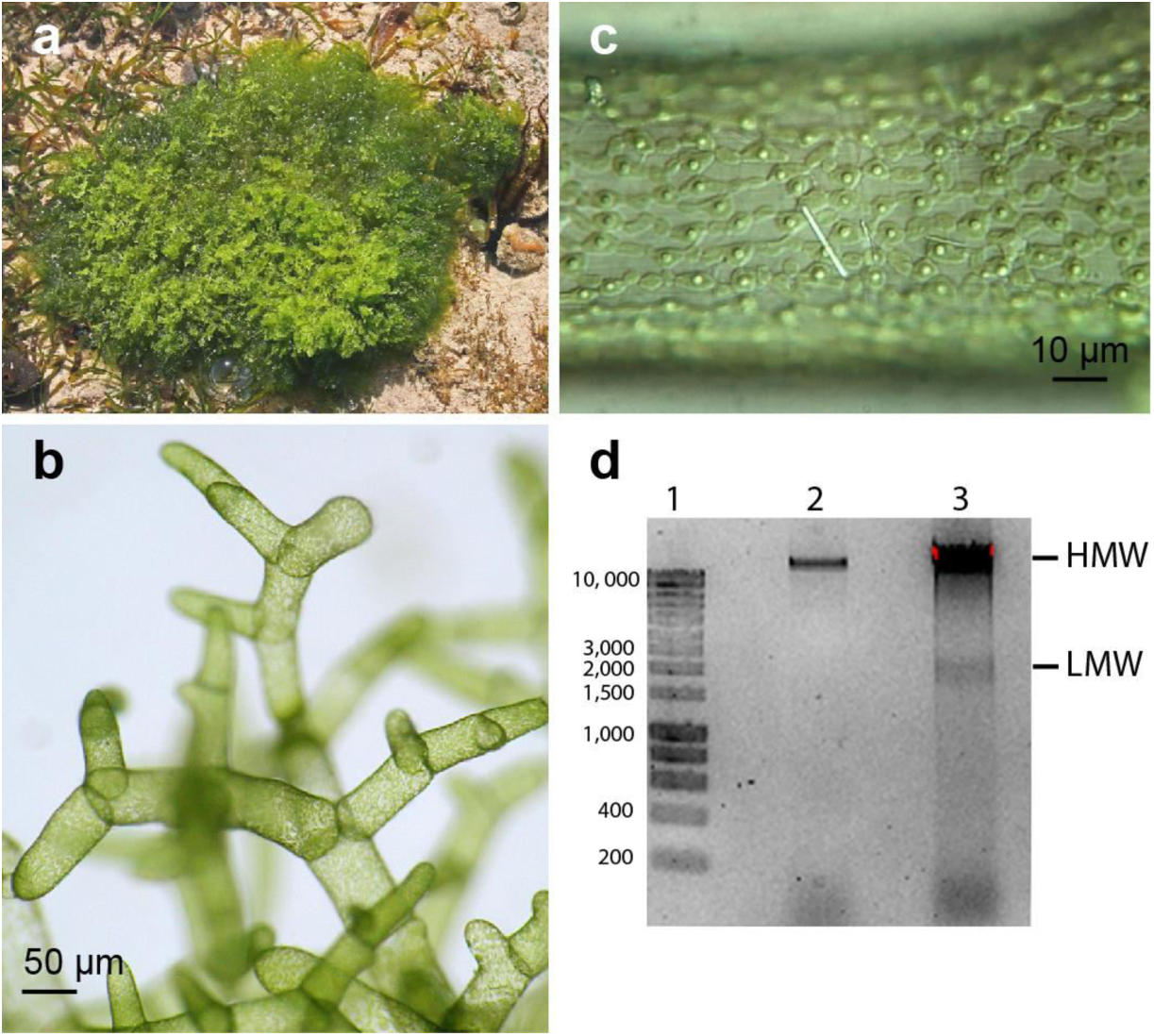
Boodlea composita. **a.** Specimen in natural environment. **b.** Detail of branching cells. **c.** Detail of chloroplasts, each containing a single pyrenoid, and forming a parietal network (the white line is a calcium oxalate crystal). **d.** Native agarose gel comparing genomic DNA of *Bryopsis plumosa* (Bryopsidales) and *Boodlea composita* (Cladophorales). Lane 1: 1-kb ladder, sizes in bp; lane 2: *B. plumosa*; lane 3: *B. composita*. High molecular weight (HMW) and low molecular weight (LMW) DNA bands of *B. composita* are indicated.

Here, we describe intriguing features of the plastid genome of Cladophorales, focusing on *Boodlea composita*. Through the integration of different DNA sequencing methods, combined with RNA sequencing, we found that chloroplast protein-coding genes are highly expressed and are encoded on 1-7 kb linear single-stranded DNA molecules. Due to the wide-spread presence of inverted repeats, these molecules fold into a hairpin configuration. A chloroplast genome that is composed only of linear DNA molecules is unprecedented among eukaryotes and highlights unexpected variation in plastid genome architecture.

## Results and Discussion

### DNA and RNAseq data

Our reconstruction of the chloroplast genome of *Boodlea composita* is based on different high-throughput DNA sequencing methods (Supplementary Fig. 1, Materials and Methods). The choice of short read DNA sequencing of a chloroplast-enriched fraction using Roche 454 technology was based on comparable sequencing approaches in other plants and algae that successfully resulted in assembly of chloroplast genomes ^30, 38^. To overcome possible assembly artefacts in a hypothetical scenario of an inflated chloroplast genome bloated by repetitive elements, long-read sequencing of the High Molecular Weight (HMW) DNA fraction using Pacific Biosciences Single-Molecule Real-Time (SMRT) method was applied, while long read sequencing of the LMW DNA fraction allowed characterization of the previously observed plasmids in the chloroplast^34, 35^. To allow comparison of the results of *Boodlea* with other species of Cladophorales, we generated additional DNA sequence data from nine other species using Illumina HiSeq 2000 technology. Finally, two deep-coverage RNA-seq libraries, a total-RNA library and a mRNA library enriched for nuclear transcripts, were generated to confirm the transcription of genes, and to inform whether genes are nuclear versus plastid encoded.

### A prodigious chloroplast genome with reduced gene set

Assembly of the chloroplast-enriched DNA reads generated using Roche 454 technology did not result in a typical circular chloroplast genome. Instead, 21 chloroplast protein-coding genes were found on 58 short contigs (1,203-5,426 bp): *atpA*, *atpB*, *atpH*, *atpI*, *petA*, *petB*, *petD*, *psaA*, *psaB*, *psaC*, *psbA*, *psbB*, *psbC*, *psbD*, *psbE*, *psbF*, *psbJ*, *psbK*, *psbL, psbT* and *rbcL*. All but the *rbcL* gene code for components of the major thylakoid transmembrane protein complexes (ATP synthase, cytochrome b6f, and photosystems I and II). The contigs contained inverted repeats at their termini (Fig. 2, group E contig) and, despite high coverage by sequence reads, they could not be extended by iterative contig extension. Sequence similarity searches and a metagenomic binning approach (distribution analysis of 4-mers) demonstrated that the inverted repeats were also found on contigs with no sequence similarity to known proteins, raising the number of contigs of chloroplast origin to 136. These contigs are further referred to as “chloroplast 454 contigs”. The length distribution of the chloroplast 454 contigs was consistent with the size of the LMW DNA fraction as estimated by agarose gel electrophoresis of *Boodlea* genomic DNA (Fig. 1d, Supplementary Fig. 14).

**Figure 2.**
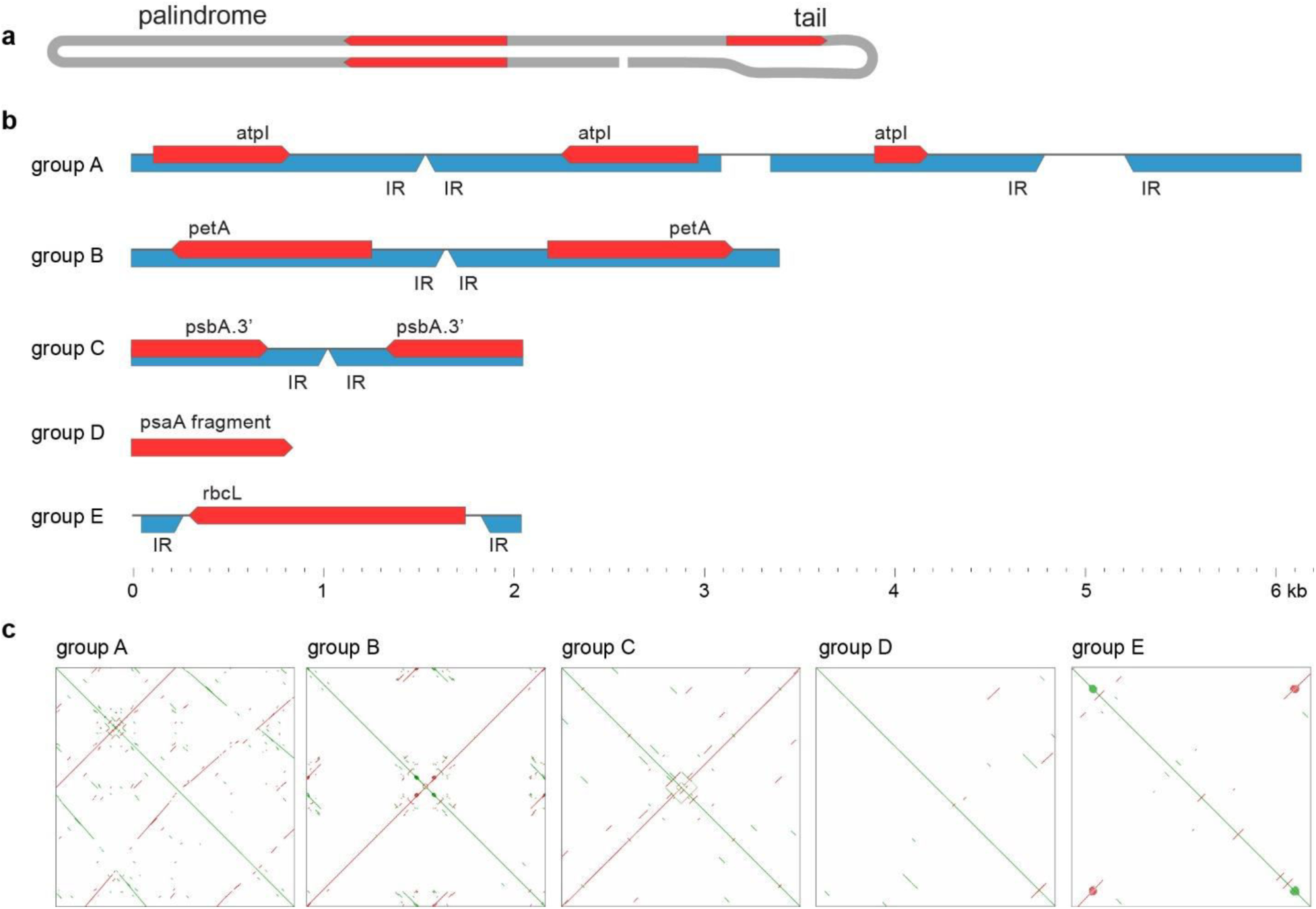
Schematic representation of assembled *Boodlea* chloroplast DNA contigs. **a.** Schematic representation of one of the predicted native conformation of chloroplast hairpins plasmids (with a near-perfect palindromic region and a less conserved tail). Red arrows represent CDSs. **b.** Schematic representation of the assembled *Boodlea* chloroplast DNA contigs; red arrows represent CDSs, blue arrows represent major inverted repeats; scale bar at the bottom indicates length of the contigs/reads. Group A: The first half of the read is a perfect palindrome, containing the two inverted CDSs; the second half of the read (tail) is less conserved, but similar to the first half of the read. Group B: Palindromic sequences with full-length CDSs in opposite orientations, resembling the first half of group A read. Group C: palindromic sequences with fragments of CDS. Group D: short reads with fragments of CDSs that lack extensive repetitive elements. Group E: Full-length CDSs delimited by inverted repeats. Similar to group E contigs and reads, the remaining 113 chloroplast 454 contigs lacking full-length CDSs are delimited by inverted repeats. **c.** Dotplots of the five groups, showing the abundance of repetitive elements. Each dotplot was generated by aligning the contig/read with itself. Green lines indicate similar sequences; red lines indicate sequences similar to the respective reverse complements.

The failure to assemble a circular chloroplast genome might be due to repetitive elements that impair the performance of short-read assemblers^39^. Inflated chloroplast genome bloated by repetitive elements have been documented in several green algae ^40, 41, 42^. To overcome assembly artefacts and close putative gaps in the chloroplast 454 contigs, we applied Single-Molecule Real-Time (SMRT) sequencing (Pacific Biosciences) to the HMW and LMW DNA fractions. Only 22 HMW DNA reads (ca. 0.044 %) harboured protein-coding genes commonly present in chloroplast genomes of Archaeplastida (Fig. 3). All, but three of these genes (which likely correspond to carry-over LMW DNA), contained introns, were absent in the chloroplast 454 contigs, and revealed a high ratio between mapped mRNA and total-RNA reads, altogether suggesting that they are encoded in the nucleus (Supplementary Fig. 2). Conversely, 22 chloroplast genes (that is, the 21 protein-coding genes identified in chloroplast 454 contigs as well as the 16S rRNA gene) were found in the LMW DNA reads (Fig. 3). An orthology-guided assembly, where the chloroplast 454 contigs harbouring protein-coding genes guided the assembly of LMW DNA reads with sequence similarity to chloroplast genes, resulted in 34 contigs between 1,179 and 6,925 bp in length, henceforth referred to as “chloroplast genome” (Fig. 4, Supplementary Table 1).

**Figure 3.**
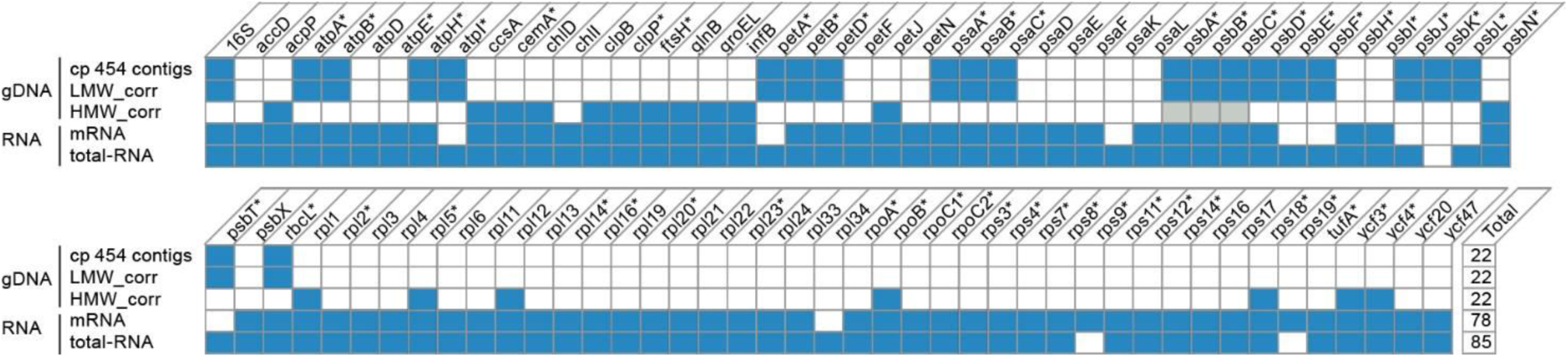
*Boodlea* genes having orthologs in the chloroplast of other Archaeplastida and their distribution over the sequenced DNA/RNA. gDNA (genomic DNA): chloroplast (cp) 454 contigs, HMW and LMW corrected reads; RNA: mRNA and total-RNA assemblies. Asterisks (*) indicate "core" chloroplast genes, i.e. protein-coding genes conserved between chloroplast genomes of Chlorophyta (see Methods). The following 9 "core" chloroplast genes were not found in any of the *Boodlea* libraries sequenced: *atpF, petG, petL, psaJ, psbM, psbZ, rpl36, rps2* and *ycf1.* Grey cells are putative carry-over contaminants: LMW DNA reads contaminants as suggested by the ratios of HMW to LMW DNA reads and mRNA to total-RNA reads (Supplementary Fig. 2).

Several contigs of the *Boodlea* chloroplast genome display long imperfect palindromic sequences that include full-length coding sequences (CDSs), and a less conserved tail region (Fig. 2). The remaining contigs have similar palindromic structures but appear to be not completely assembled. Such palindromes allow regions of the single-stranded LMW DNA molecules to fold into hairpin-like secondary structures. Additional smaller inverted repeats were identified in many of the contigs, which may result in more complex secondary structures (Fig. 4).

**Figure 4.**
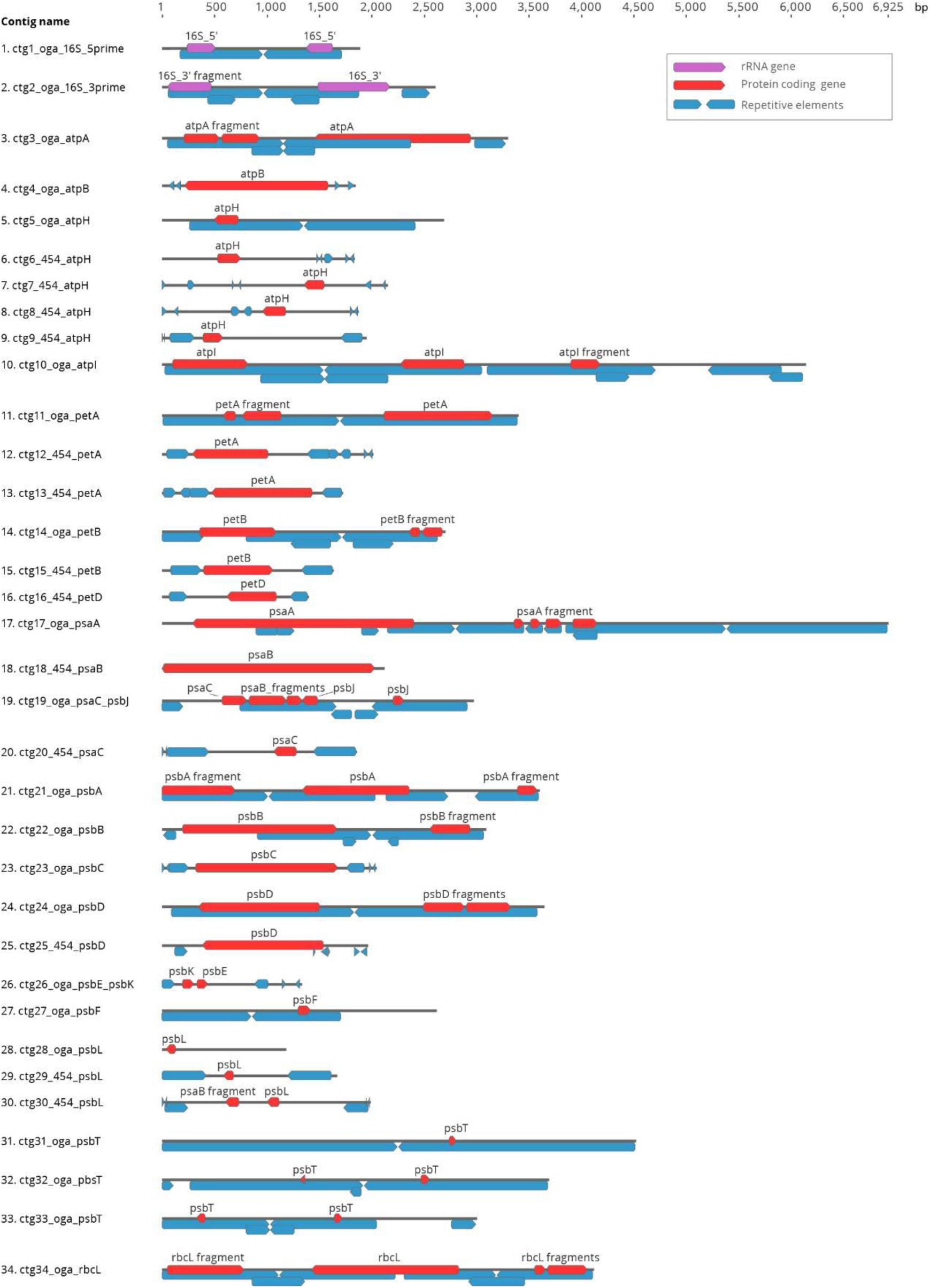
Schematic representation of *Boodlea* chloroplast genome. Purple arrows indicate rRNA genes, red arrows indicate CDSs of protein-coding genes, and blue arrows indicate repetitive elements. For each contig, repetitive elements with similar length indicate similar sequences. Oga: contig obtained by orthology-guided assembly. 454: chloroplast 454 contig.

Chloroplast 454 contigs could not be scaffolded with long HMW DNA reads, nor did an hybrid assembly between chloroplast 454 contigs and long HMW DNA reads generate a circular chloroplast genome (Supplementary Figs. S1, Materials and Methods). The LMW DNA reads are concordant and consistent with the imperfect palindromic sequences of the assembled chloroplast genome, indicating that the palindromes are not a result of assembly artefacts (Fig. 4, Supplementary Table 1). As a consequence, we conclude that the chloroplast genome is not a single large molecule but it is instead fragmented in several molecules in the LMW DNA.

A chloroplast genome that is entirely fragmented into plasmids is in line with earlier observations of abundant LMW DNA in chloroplasts of several species of Cladophorales^34^. The hairpin configuration of the plasmids derived from our sequence data corresponds with earlier data based on electron microscopy, endo-and exonuclease digestion experiments, acridine orange staining, and denaturing gel electrophoresis^34, 37^. Fluorescence *in situ* hybridization, and Southern blot hybridisation indicated that these plasmids were present within the chloroplast only^35^, supporting the congruence between chloroplast 454 contigs and sequences from the LMW fraction (Fig. 3).

The chloroplast genome contigs of *Boodlea* feature an exceptionally high GC-content, ranging from 54 to 60 % in the gene-containing contigs (average 57%) (Supplementary Table 1). These values are concordant with the high density of the LMW fraction observed in CsCl/ bisbenzimide gradients^34^, and also with sequence data from cloned plasmids of *Ernodesmis* (51- 59% GC)^36^. Plastid genomes are generally AT-rich, and in green algal species GC-content typically ranges between 26 and 43%^12, 43^. GC-rich plastid genomes are very rare, but higher values have been reported for the trebouxiophycean green algae *Coccomyxa subellipsoidea*, *Paradoxia multiseta* (both 51% GC), and Trebouxiophyceae sp. MX-AZ01 (58%)^29^. These species, however, feature standard plastid genome architectures.

The size of the *Boodlea* chloroplast genome could not be estimated by inspection of k-mer frequency distributions of the reads in the 454 library nor from those of the uncorrected and corrected LMW DNA reads^44^. Histograms of k-mer frequency distributions revealed several small peaks, indicating a heterogeneous population of molecules present in different stoichiometries, and the signal to noise ratio was too small to make a comfortable estimation of the sizes (Supplementary Fig. 3). The cumulative length of the 34 *Boodlea* chloroplast genome contigs is 91 kb (Supplementary table 3). However, if we would consider the large and heterogeneous population of LMW DNA reads bearing no similarity to protein-coding genes (“empty” plasmids, see below) as part of the chloroplast genome, its size could be regarded as much larger.

The largest known circular-mapping chloroplast genomes have been documented for two clades of green algae, the Chlorophyceae and the Ulvophyceae. Inflation of the 521-kb chloroplast genome of *Floydiella terrestris* (Chlorophyceae) resulted mainly from the proliferation of dispersed, heterogeneous repeats (>30 bp) in intergenic regions, representing more than half of the genome length^42^. Intergenic regions of the *Volvox carteri* (Chlorophyceae) chloroplast genome, instead, are populated with short palindromic repeats (average size of 50 bp) that constitute ca. 64% of the predicted 525-kb genome^40^. The mechanisms by which such palindromic selfish DNA spread throughout the *Volvox* chloroplast genome is not clear, but the presence of a reverse transcriptase and endonuclease may point toward retrotranscription^40, 45^. For *Acetabularia acetabulum* (Ulvophyceae), the chloroplast genome was sequenced only partially and its size was estimated to exceed 1 Mb; it has exceptionally long intergenic regions and features long repetitive elements (>10 kb) arranged in tandem^41, 46^. The *Boodlea* chloroplast genome is rich as well in non-coding DNA, constituting 92.2% of the 136 chloroplast 454 contigs and 72.8% of the assembled chloroplast genome, comparable to that in inflated chloroplast genomes of other green algae (*Floydiella terrestris,* 82.1%; *Volvox carteri,* ca. 80%; *Acetabularia acetabulum,* ca. 87% of the sequenced chloroplast genome)^40, 41^.

The non-coding DNA regions (ncDNA) of the hairpins showed high sequence similarity among one another. Within the ncDNA, we identified six conserved motifs, 20 to 35 bp in length, which lack similarity to known regulatory elements (Fig. 6). Motifs 1, 2 and 5 were always present upstream of the start codon of the chloroplast genes, occasionally in more than one copy. Although their distances from the start codon were variable, their orientations relative to the gene were conserved, indicating a potential function as a regulatory element of gene expression and/or replication of the hairpin plasmids. These motifs were also present in 1,966 (ca. 1.8 %) LMW DNA reads lacking genes. This observation supports earlier findings of abundant non-coding LMW DNA molecules in the Cladophorales^34, 36^. In contrast, a very small fraction of the HMW DNA reads (15 corrected reads) displayed the same ncDNA motifs and these were present exclusively on long terminal repeat retrotransposons (RT-LTRs) (Supplementary Fig. 4, Supplementary Note 1). RT-LTRs were also abundant in the 454 contigs (Supplementary Fig. S16). The abundance of RT-LTRs in the 454 contigs and the presence of ncDNA motifs in both the *Boodlea* chloroplast genome and nuclear RT-LTRs is suggestive of DNA transfer between the nucleus and chloroplast and may allude to the origin of the hairpin plasmids. Hypothetically, an invasion of nuclear RT-LTRs in the chloroplast genome may have resulted in an expansion of the chloroplast genome and its subsequent fragmentation into hairpin plasmids during replication. Chloroplast genome fragmentation could be caused by recombination between repetitive elements and displacement of the palindromic sequences from the lagging strand during the chloroplast genome replication^47, 48^, and it is consistent with the expectation that recombination and cleavage of repetitive DNA will produce a heterogeneous population of molecules, as observed in dinoflagellates plastid genomes^14^, and in the *Boodlea* LMW DNA.

### A fragmented chloroplast genome is a common feature of Cladophorales

DNA sequence data were obtained from 9 additional Cladophorales species, representing the main lineages of the order: *Chaetomorpha aerea*, *Cladophora albida*, *C. socialis*, *C. vadorum*, *Dictyosphaeria cavernosa*, *Pithophora* sp., *Siphonocladus tropicus*, *Struvea elegans*, and *Valonia utricularis* (Supplementary Table 7). Although comparable sequencing approaches resulted in the assembly of circular chloroplast genomes for other algae, including green seaweeds^49, 50, 51^, only short chloroplast contigs (200-8,000 bp) were assembled from these libraries, similar to *Boodlea composita*. Interestingly, a similar set of chloroplast genes was identified in all sequenced Cladophorales species. In contrast to the genes found in the *Boodlea* hairpin plasmids, however, most of the chloroplast genes identified in the additional Cladophorales libraries were fragmented, probably due to sequencing of the shorter Illumina reads (Supplementary Table 9). These findings support the idea that fragmentation of the chloroplast genome occurred before or early in the evolution of the Cladophorales.

### Highly divergent chloroplast genes are transcribed

The 21 chloroplast protein-coding genes of *Boodlea* and the other species of Cladophorales display extremely high sequence divergence compared to orthologous genes in other photosynthetic organisms (Fig. 7b). A maximum likelihood phylogenetic tree based on a concatenated alignment of 19 chloroplast genes from Archaeplastida and Cyanobacteria species is presented in Fig. 7a. Despite their high divergence, the Cladophorales sequences formed a monophyletic group within the core Chlorophyta, sister to the Trentepohliales (Supplementary Fig. 5), a position that is supported by phylogenetic analyses of nuclear genes^52, 53^. The high sequence divergence of chloroplast genes in the Cladophorales supports the notion that organellar genomes with extremely derived architectures, including those of peridin-containing dinoflagellates, also tend to fall at the extreme ends of the range observed at the mutation rate (or gene sequence divergence) level^11, 54^.

**Figure 5.**
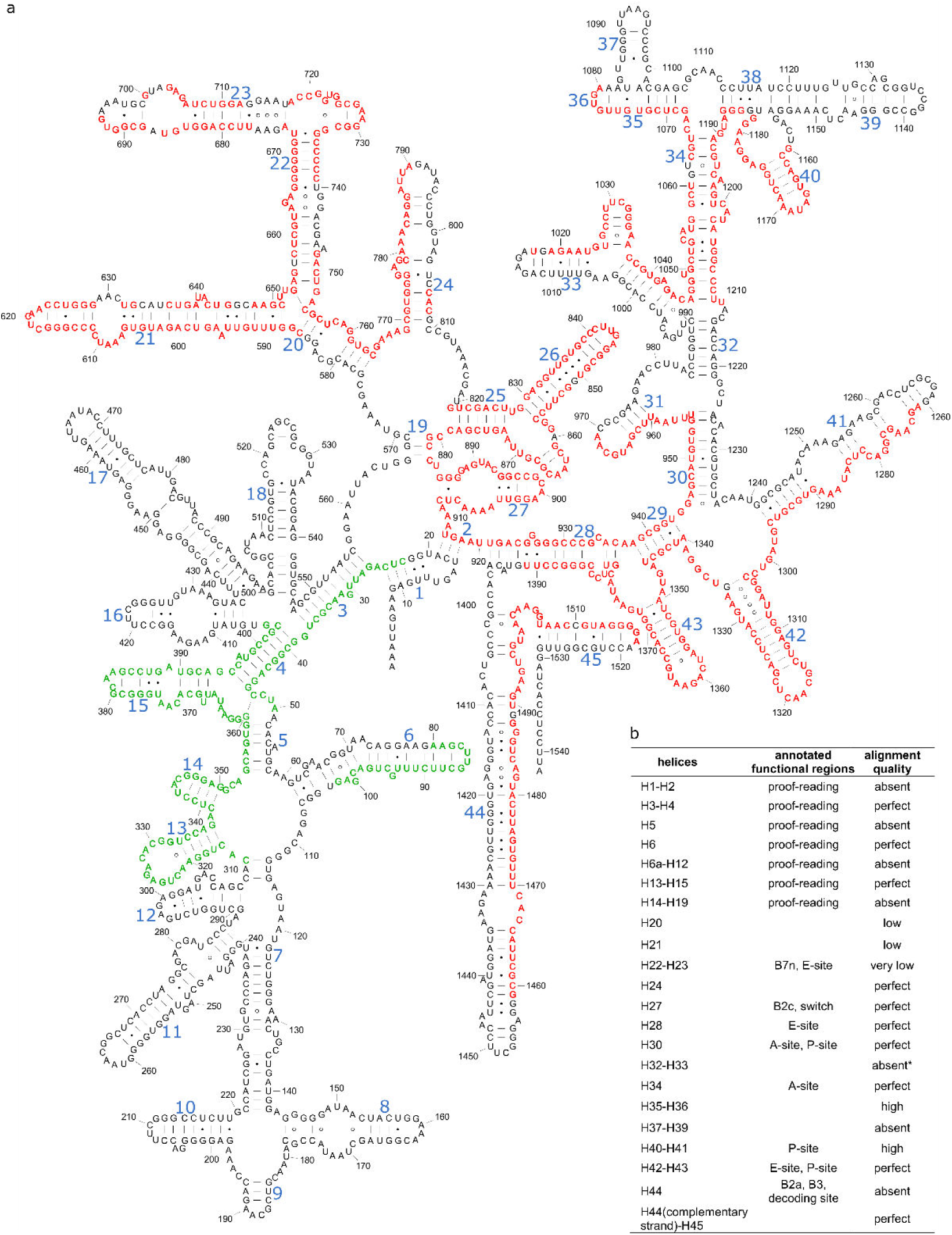
*Boodlea* chloroplast 16S rRNA secondary structure. **a.** *Boodlea* chloroplast 16S rRNA secondary structure was compared with the *E. coli* 16S rRNA model [RF00177]. Residues shown in green and red on the *E*. *coli* model represent the 5’ and 3’ 16S rRNA regions coded by the two *Boodlea* 16S rRNA hairpin plasmids. Residues in black are absent in *Boodlea* 16S rRNA. Blue numbers indicate secondary structure helices in the 16S rRNA model ^102^. **b.** Comparison between *Boodlea* and *E. coli* 16S rRNA annotated functional regions. Quality of the alignment was assessed based on the predicted posterior probability (in percentage) of each aligned region: very low < 25%; low between 25-50%; high between 50-95%; and perfect > 95%.

**Figure 6.**
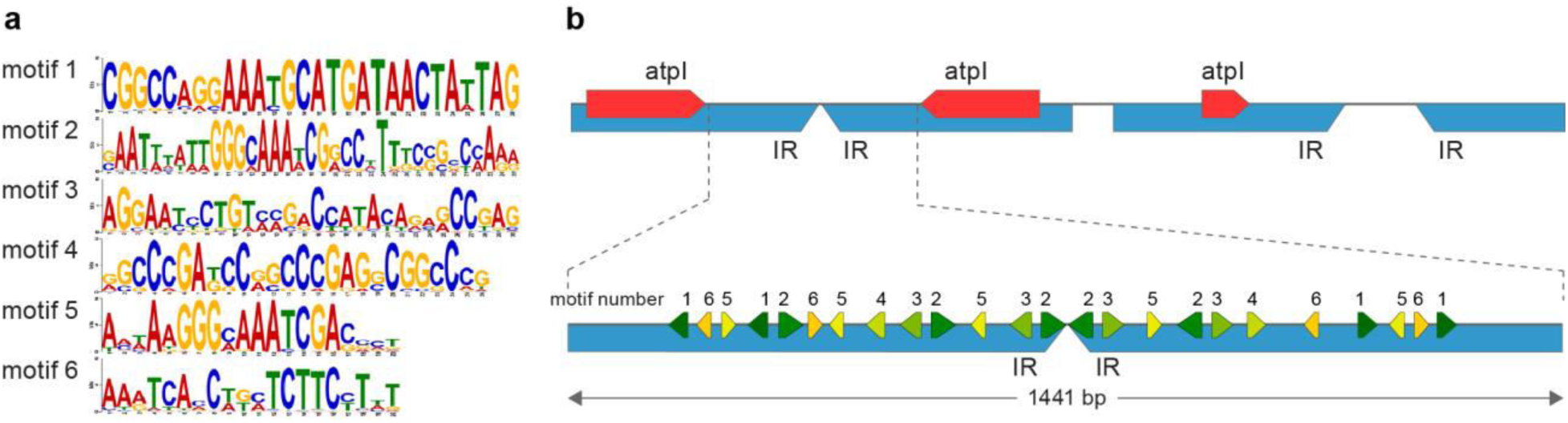
Conserved non-coding motifs in *Boodlea* LMW DNA. **a.** Sequence logos of the conserved motifs predicted in the *Boodlea* chloroplast genome. **b.** Schematic representation of the distribution of the motifs in the 1,441 bp ncDNA region from the *atpI* group A read used for the identification of additional chloroplast reads in the LMW DNA library. Motifs with conserved orientation relative to the downstream genes are represented by green arrows, while motifs without conserved orientation to the downstream genes are represented by yellow arrows. CDSs are represented by red arrows, inverted repeats are represented by blue arrows.

**Figure 7.**
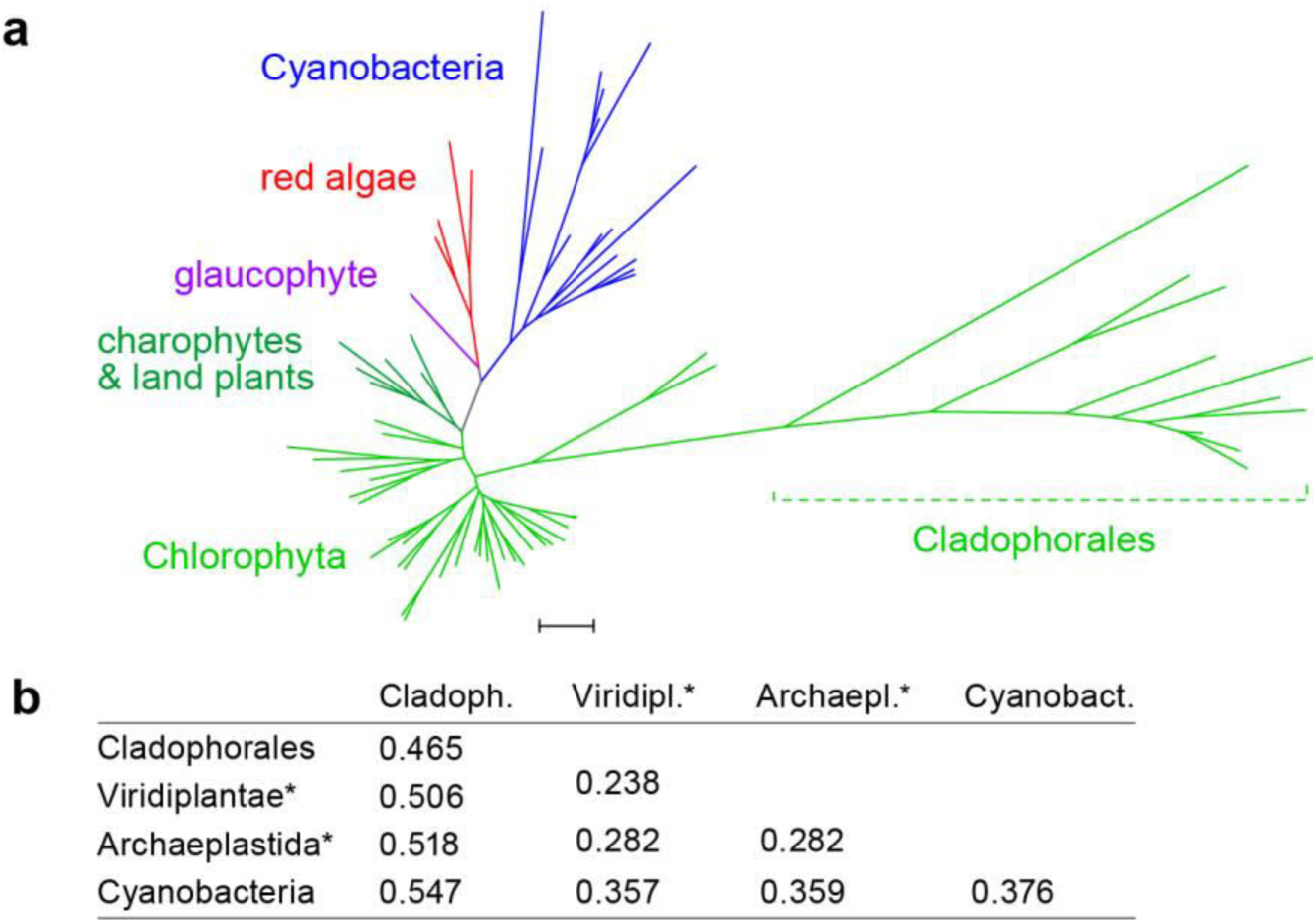
Phylogeny of Archaeplastida and Cyanobacteria based on 19 chloroplast genes, and corresponding distance matrix illustrating the large sequence divergence of Cladophorales chloroplast genes. **a.** Maximum likelihood phylogenetic tree inferred from a concatenated amino acid alignment of 19 chloroplast genes. The phylogeny with taxon names and bootstrap support values is shown in Supplementary Fig. 4. The scale represent 0,3 substitution per amino acid position. **b.** Maximum pairwise amino acid sequence distances of the chloroplast genes within and between clades (* excluded Cladophorales).

For some *Boodlea* chloroplast genes, the identification of start and stop codons was uncertain and a non-canonical genetic code was identified (Supplementary Fig. 6). The canonical stop codon UGA was found 11 times internally in six genes (*petA*, *psaA*, *psaB*, *psaC*, *psbC* and *rbcL*), but was also present as a genuine termination codon in several genes, *petA* and *psaA* included. At seven of these 11 positions, the corresponding amino acid residue in orthologous genes was conserved (i.e. present in more than 75% of the taxa in the alignment), but different amino acids were observed at these positions: V, Q, I and L (Supplementary Fig. 6b). For the remaining positions, the amino acid in the alignment was not conserved, and therefore the amino acid coded by the UGA codon could not be determined with certainty.

Deviations from the universal genetic code are widespread among mitochondrial genomes, and include loss of start and stop codons in some groups, including dinoflagellates^14, 55, 56^. In contrast, non-canonical genetic codes are much rarer in plastid genomes, and up to now have only been detected in the apicomplexans *Neospora caninum*^57^, *Chromera velia*^26^, and the dinoflagellate *Lepidodinium chlorophorum*^58^. In genomes of primary plastids, a non-canonical genetic code is unprecedented.

Dual meaning of UGA as both stop and sense codons has recently been reported from a number of unrelated protists^59, 60, 61, 62, 63^. While in *Saccharomyces cerevisiae* the tetranucleotide UGA-C allows increased incorporation of the near-cognate Cys-tRNA for the UGA premature termination codon^64^, such preference was not observed in the *Boodlea* chloroplasts. Importantly, a non-canonical genetic code has also been described for Cladophorales nuclear genes, where UAG and UAA codons are reassigned to glutamine^65^, which implies two independent departures from the standard genetic code in a single organism.

Unexpectedly, we found that the 16S rRNA gene in the *Boodlea* chloroplast genome is split across two distinct hairpin plasmids and that its size is much smaller compared to its algal and bacterial homologs and (Figs 4, 5). Fragmentation of rRNA genes has been observed in organellar genomes, including Apicomplexa, dinoflagellates, and many green algae^13^. In general, fragmentation of protein-coding and rRNA genes is more common in mitochondrial genomes than in plastid genomes^21, 26, 68, 69^. Despite considerable effort, we could not detect the 23S rRNA gene nor the 5S rRNA gene.

The transcription of the aberrant chloroplast genes was confirmed using RNAseq, and is concordant with previous results of Northern blots^36^. Transcripts of 21 chloroplast genes (that is, 20 protein-coding genes as well as the 16S rRNA gene) were identical to the genes encoded by the chloroplast 454 contigs (Figs. 3, 8, Supplementary Figs 7-12), providing evidence for the absence of RNA editing and corroborating the use of a non-canonical genetic code (Supplementary Figs 7-12). Lack of RNA editing was also evidenced for the 11 internal occurrences of UGA (Supplementary Figs 7-12). This observation, in combination with conservation of the sequence after the UGA codon, serves as evidence that it is not a termination codon but an alternative code. The high total-RNA to mRNA ratio observed for reads that mapped to the chloroplast 454 contigs confirmed that these genes were not transcribed in the nucleus (Supplementary Fig. 17). All coding sequences of the same protein-coding genes found on different contigs of the *Boodlea* chloroplast genome were expressed, despite minor differences in their nucleotide sequences (Supplementary Table 1).

**Figure 8.**
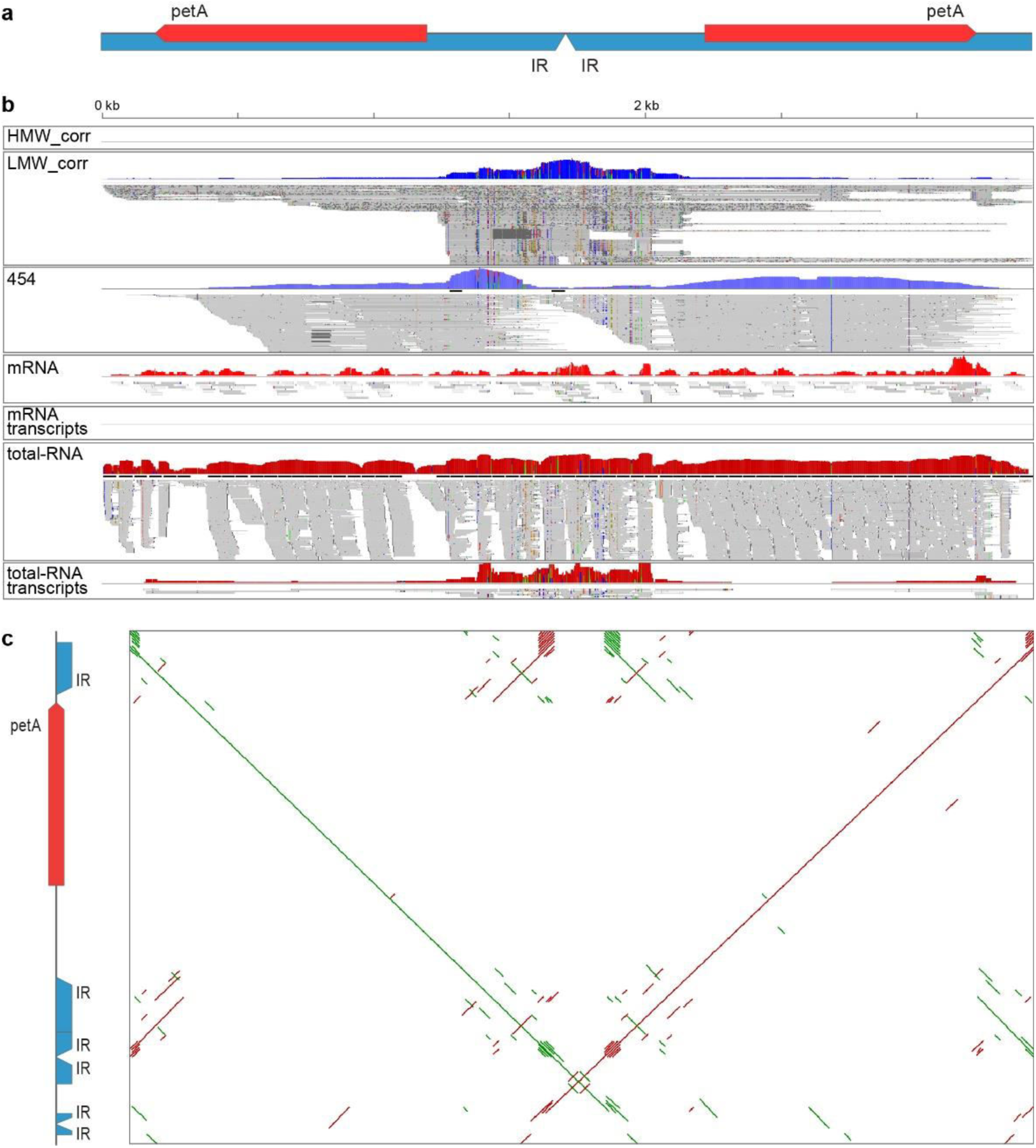
LMW DNA reads containing chloroplast genes are expressed, enriched in the total-RNA fraction and congruent to the respective chloroplast 454 contigs. **a.** Representation of *petA* LMW DNA read (3,398 bp), a representative of group B contigs and reads (see Fig. 2). The red arrows indicate CDSs, the blue arrows indicate inverted repeats. **b.** Corresponding Genome Browser track, from top to bottom: corrected HMW DNA coverage [0], corrected LMW DNA read coverage [range 0-541], 454 read coverage [range 0-567], mRNA library read coverage [range 0-17], assembled mRNA transcripts mapped [0], total-RNA library read coverage [range 0-7,936], and assembled total-RNA transcripts mapped [range 0-17]. **c.** Dotplot showing congruence between *petA* LMW DNA read (x axis) and the corresponding *petA*-containing chloroplast 454 contig (y axis, 2,012 bp). Green lines indicate similar sequences; red lines indicate sequences similar to the respective reverse complements.

Additional transcripts of 66 genes that have been located in the chloroplast in other Archaeplastida were identified (Fig. 3). Although their subcellular origin was not determined experimentally, they are probably all nuclear-encoded, based on high mRNA to total-RNA reads ratio and their presence on High Molecular Weight (HMW) DNA reads.

### Conclusions

We collected several lines of evidence indicating that *Boodlea composita* lacks a typical large circular chloroplast genome. The chloroplast genome is instead fragmented into multiple linear hairpin plasmids, and has a highly reduced gene repertoire compared to other chloroplast genomes. Thirty-four hairpin plasmids were identified, harbouring 21 protein-coding genes and the 16S rRNA gene, which are highly divergent in sequence compared to orthologs in other algae. The exact set of *Boodlea* chloroplast genes remains elusive, but at least 19 genes coding for chloroplast products appear to be nuclear-encoded, of which nine are always chloroplast-encoded in related green algae (Fig. 3). This suggests that fragmentation of a conventional chloroplast genome in the Cladophorales has been accompanied with an elevated transfer of genes to the nucleus, similarly to the situation in peridinin-containing dinoflagellates^14^, with plastid genomes encoding about 12 genes or less^14, 70^. Notably, the two distantly related algal groups have converged on a very similar gene distribution: chloroplast genes code only for the subunits of photosynthetic complexes (and also for Rubisco in *Boodlea*), whereas the expression machinery is fully nucleus-encoded (Table 1). Other nonstandard chloroplast genome architectures have recently been observed, such as a monomeric linear chromosome in the alveolate microalga *Chromera velia*^26^ and three circular chromosomes in the green alga *Koshicola spirodelophila*^15^, but these represent relatively small deviations from the paradigm, when compared to the chloroplast genome of the Cladophorales. The highly fragmented chloroplast genome in the Cladophorales is wholly unprecedented and will be of significance to understanding processes driving organellar genome fragmentation and gene reduction, endosymbiotic gene transfer, and the minimal functional chloroplast gene set.

### Methods

#### Algal material and chloroplast isolation

A clonal culture of *Boodlea composita* (FL1110) is maintained in the algal culture collection of the Phycology Research Group, Ghent University. The specimen was grown in enriched sterilized natural seawater^71^ at 22°C under 12:12 (light:dark) cool white fluorescent light at 60 µmol photons m^-2^ s^-1^. A chloroplast-enriched fraction was obtained following the protocol of Palmer et al.^72^.

#### Genomic DNA extraction, quantification and sequencing

Total genomic DNA from fresh *Boodlea* cultures was isolated by using a modified CTAB extraction protocol^73^. HMW and LMW DNA bands were size-selected using a BluePippin^TM^ system (Sage Science, USA). The HMW DNA band was isolated with a cut-off range of 10 kb to 50 kb, while the LMW DNA band was isolated with a cut-off range of 1.5 kb to 2.5 kb. The quantity, quality and integrity of the extracted DNA were assessed with Qubit (ThermoFisher Scientific, USA), Nanodrop spectrophotometer (ThermoFisher Scientific), and Bioanalyzer 2100 (Agilent Technologies, USA).

DNA from the chloroplast-enriched fraction was sequenced with Roche 454 GS FLX at GATC Biotech, Germany. The HMW and LMW DNA fractions were sequenced on two SMRT cells on a PacBio RS II (VIB Nucleomics Core facilities, Leuven, Belgium) using PacBio P5 polymerase and C3 chemistry combination (P5-C3). For the HMW DNA fraction, a 20-kb SMRT-bell library was constructed, while for the LMW DNA fraction, a 2-kb SMRT-bell library was constructed.

#### Chloroplast DNA assembly and annotation

Quality of the reads from the 454 library was assessed with FastQC v.0.10.1 (<u>http://www.bioinformatics.babraham.ac.uk</u>, last accessed March 01, 2017) (Supplementary Fig. 13, Supplementary table 2). Low-quality reads (average Phred quality score below 20) were discarded and low-quality 3’ ends of the reads were trimmed with Fastxv.0.0.13 (https://github.com/agordon/fastx_toolkit, last accessed March 01, 2017). After trimming, reads shorter than 50 bp were discarded. *De novo* assembly of the trimmed reads was performed with MIRA v. 4.0rc5^74^. The assembly resulted in 3,735 contigs, which will be further referred to as “454 contigs” (Supplementary table 3). Length distribution of the 454 contigs is reported in Supplementary Fig. 13.

After the assembly, putative chloroplast contigs were identified by comparing their translated sequences against the NCBI non-redundant protein database using BLAST 2.2.29+^75^, resulting in the identification of 58 contigs harbouring fragments or full-length chloroplast genes by sequence similarity search. These contigs had long stretches of conserved repetitive sequences at their 5’ and 3’ extremities. The conserved inverted repeats were used in a sequence similarity search with high stringency (high mismatch cost, high cost for gap opening and gap extension, long minimal word-size) against the 454 contigs to identify 18 additional contigs of putative chloroplast origin. This initial set of 76 contigs had a mean coverage of 84×, ranging between 11× and 191×. 17 of the 76 contigs had internal inverted repeats, with a sudden drop in read coverage (Supplementary Fig. 15). These contigs were regarded as chimeric contigs and were cleaved at the sites of coverage drop, raising the number of contigs of chloroplast origin to 89.

Additional chloroplast contigs without similarity to protein-coding genes were identified by metagenomic binning (distribution analysis of 4-mers) with MyCC^76^, resulting in 21 clusters of 454 contigs (Supplementary Fig. 16). The initial set of 89 chloroplast 454 contigs was present in three neighbouring clusters: Cluster 14, Cluster 17 and Cluster 21. These clusters contained 122, eight and six contigs, respectively, raising the number of identified chloroplast contigs assembled from the chloroplast-enriched fraction from 89 to 136 (“chloroplast 454 contigs” in Supplementary table 3). Of these, 71 contigs had no sequence similarity to known protein-coding genes, 36 contigs harboured full-length chloroplast genes, 36 contigs harboured fragments of chloroplast genes, and 7 contigs harboured both fragments and full-length CDSs of different chloroplast genes.

Contigs potentially coding for chloroplast tRNAs and rRNAs were identified using Infernal 1.1^77^. The chloroplast 454 contigs served as seeds for iterative contig extension with PRICE^78^. Single-end 454 reads were used as false paired-end reads with expected insert size equal to the median length of the 454 reads. 141 different combinations of parameters were tested in order to optimize the contig extension. None of the selected assemblies showed a length improvement for the initial set of chloroplast 454 contigs. The length distribution of the chloroplast 454 contigs was consistent with the size of the LMW DNA fraction as estimated by agarose gel electrophoresis of *Boodlea* genomic DNA (Fig. 1d, Supplementary Fig. 14).

Repetitive regions in the contigs were identified with ‘einverted’, ‘etandem’ and ‘palindrome’ from the EMBOSS^79^ package. Dotplots for all contigs were generated with YASS, using standard parameters^80^. Coverage of the chloroplast 454 contigs was evaluated by mapping the 454 reads, the mRNA and the total-RNA libraries to these contigs with CLC Genomics Workbench 7.0 (Qiagen) (Supplementary Fig. 17).

The chloroplast 454 contigs were used together with HMW DNA reads for two independent hybrid assemblies. First, we tried to close hypothetical gaps between the chloroplast 454 contigs with the pbahaScaffolder.py script integrated in the smrtanalysis 2.3.0 pipeline^81^. Secondly, the pre-assembled chloroplast 454 contigs were used as anchors for HMW DNA reads in a round of hybrid assembly with dbg2olc^82^. These analyses failed to close the hypothetical gaps between the short chloroplast 454 contigs and did not yield longer contigs. These results stand in stark contrast to the mitochondrial 454 contigs, where the same approaches yielded markedly longer contigs (see Supplementary Note 2).

Since the hybrid assemblies with uncorrected reads could not reconstruct a circular chloroplast genome, HMW DNA reads were further characterized after error correction. The high-noise HMW DNA reads were corrected by applying a hybrid correction with proovread 2.12^83^ using 454 reads and reads from Illumina RNA-seq libraries (see below). Corrected reads encoding chloroplast genes were identified by aligning them against a custom protein database, named Chloroprotein_db, including genes from the pico-PLAZA protein database^84^ and protein-coding genes from published green algal chloroplast genomes (Chlorophyta sensu Bremer 1985, NCBI Taxonomy id: 3041).

LMW DNA reads were self-corrected with the PBcR pipeline^85^. Since the LMW DNA size is unknown and PBcR requires an estimate of the genome size for proper read correction, six different putative genome sizes were tested (100 kb, 1 Mb, 2.24 Mb, 10 Mb, 100 Mb). The best performance in terms of number of corrected reads was obtained by the combination of "10 Mb" for the estimated genome size and the *–sensitive* flag turned on; these corrected reads were used for the downstream analysis. After error correction, the number of reads was reduced from 154,852 to 106,428 (Supplementary table 2), with a similar length distribution as the uncorrected reads library (Supplementary Fig. 13).

In order to estimate the *Boodlea* chloroplast genome size, k-mer frequency distributions were calculated with jellyfish 2.0^86^. K-mers ranging from 11 toto 47 were analysed for uncorrected and corrected LMW DNA reads, for the filtered 454 reads and for the 454 reads that could be mapped on the chloroplast 454 contigs (Supplementary Figure 3).

*De novo* genome assembly of corrected LMW DNA reads was performed with the Celera WGS assembler version 8.3rc2^87^. The resulting assembly, hereafter called the Celera Assembly, consisted of 558 contigs (Supplementary table 3). Corrected and uncorrected reads as well as assembled contigs potentially encoding chloroplast genes were identified by aligning them with BLAST 2.2.29+ against Chloroprotein_db. In order to identify additional short protein-coding genes, HMM profiles were generated from alignments of chloroplast genes present in Chloroprotein_db and used to search the 6-frame translations of 454 contigs and corrected and uncorrected HMW and LMW DNA reads with HMMer3^88^. To prevent assembly artefacts caused by repetitive elements and palindromic sequences, we also performed an orthology-guided assembly, in which the LMW DNA reads harbouring chloroplast CDSs were re-assembled together with the respective chloroplast 454 contigs. First, LMW DNA corrected reads and chloroplast 454 contigs were grouped according to their best BLAST hit. The corrected reads and contigs belonging to the same group were assembled using Geneious v. 8.1.7 (Biomatters, http://www.geneious.com/, last accessed March 01, 2017) with parameters "High Sensitivity/Medium", and each assembly (or lack of assembly) was visually screened to exclude potential chimeric contigs (e.g. palindromic corrected subreads should be collapsed in the same locus rather than being concatenated). Where possible, LMW DNA reads and chloroplast 454 contigs were assembled as group A molecules (Fig. 2). The orthology-guided assembly yielded 19 contigs, 2 belonging to group A, 13 to group B and 4 to group E (Supplementary Table 1). Two groups of reads could not be assembled into longer molecules, and for them, the corresponding chloroplast 454 contigs were retained. Eleven additional chloroplast 454 contigs were retained, since they were not congruent with the LMW DNA reads and could not be included in the assembly. This resulted in a total of 32 contigs containing chloroplast protein-coding genes, which together with the two later identified Group B contigs encoding the 16S rRNA gene, are regarded as the *Boodlea* chloroplast genome contigs (Fig. 4, Supplementary table 3).

Protein-coding genes in the *Boodlea* chloroplast genome contigs were identified with a sequence similarity search against the NCBI non-redundant protein database with BLAST 2.2.29+. Their annotation was manually refined in Geneious and Artemis 16.0.0^89^ based on the BLAST search results. rRNAs were identified using Infernal 1.1^77^. Repetitive elements were mapped on the *Boodlea* chloroplast genome by aligning the contigs with themselves using BLAST 2.2.29+. Non-coding RNAs were identified with infernal 1.1 (cut-off value 10^−5^). Conserved motifs were predicted with MEME suite^90^, and the discovered motifs were clustered with RSAT^91^. The motifs were compared with the JASPAR-2016^92^ database using TOMTOM^93^ (p-value cut-off 10^−3^).

*Boodlea* chloroplast genome coverage was evaluated by mapping the 454 reads with gsnap v.2016-04-04^94^. Corrected and uncorrected LMW DNA subreads and chloroplast 454 contigs resulting from the MIRA assembly were mapped against the *Boodlea* chloroplast genome with gmap v. 2014-12-06^94^ using the *–nosplicing* flag. Due to the high number of repetitive sequences in LMW DNA reads and 454 contigs, the resulting annotated *Boodlea* chloroplast genome was carefully inspected in order to exclude sequencing and assembly artefacts.

Completeness of the chloroplast genome was evaluated by comparing the annotated chloroplast genes to a set of 60 "core" chloroplast protein-coding genes, defined as protein-coding genes conserved among the chloroplast genomes of the following representative species of Chlorophyta: *Bryopsis plumosa*, *Chlamydomonas reinhardtii*, *Chlorella vulgaris*, *Coccomyxa subellipsoidea*, *Gonium pectorale*, *Leptosira terrestris*, *Nephroselmis olivacea*, *Oltmannsiellopsis viridis*, *Parachlorella kessleri*, and *Pseudendoclonium akinetum*, and the streptophyte *Mesostigma viride* (Fig. 3).

#### RNA extraction, quantification and sequencing

Total RNA was isolated using a modified CTAB extraction protocol^95^. RNA quality and quantity were assessed with Qubit and Nanodrop spectrophotomete, and RNA integrity was assessed with a Bioanalyzer 2100. Two cDNA libraries for NextSeq sequencing were generated using TruSeq^TM^ Stranded RNA sample preparation kit (Illumina, USA): one library enriched in poly(A) mRNA due to oligo-(dT) retrotranscription and one total RNA library depleted in rRNAs with Ribo-Zero Plant kit (Epicentre, USA). The two libraries were sequenced on one lane of Illumina NextSeq 500 Medium platform at 2x76 bp by VIB Nucleomics Core facilities (Leuven, Belgium) (Supplementary Table 4).

#### Transcriptome assembly and annotation

Quality of the reads from the two RNA-seq libraries was assessed with FastQC. Low-quality reads (average Phred quality score below 20) were discarded and low-quality 3’ ends of the reads were trimmed with Fastx. After trimming, reads shorter than 30 bp were discarded. Read normalization and *de novo* assembly of the libraries were performed with Trinity 2.0.4^96^. The resulting contigs (hereafter, transcripts) were compared using sequence similarity searches against the NCBI non-redundant protein database using Tera-BLAST(tm) DeCypher (Active Motif, USA). Taxonomic profiling of the transcripts was performed using the following protocol: for each transcript, sequence similarity searches were combined with the NCBI Taxonomy information of the top ten BLAST Hits in order to discriminate between eukaryotic and bacterial transcripts or transcripts lacking similarity to known protein-coding genes (Supplementary Table 5, Supplementary Fig. 18). Transcripts classified as "eukaryotic" were further examined to assess transcriptome completeness and to identify chloroplast transcripts. These transcripts were analysed using Tera-BLAST^TM^ DeCypher against Chloroprotein_db. Transcriptome completeness was evaluated with a custom Perl script that compared gene families identified in the *Boodlea* transcriptome to a set of 1816 "core" gene families shared between Chlorophyta genomes present in pico-PLAZA 2.0^84^, following Veeckman et al. guidelines to estimate the completeness of the annotated gene space^97^ (Supplementary Table 6).

*Boodlea* chloroplast genome expression and presence of potential RNA editing were evaluated by mapping the reads from the mRNA and total-RNA libraries to the chloroplast genome contigs with gsnap, and by aligning the transcripts resulting from the *de novo* assembly of the RNA-seq libraries to the chloroplast genome contigs with gmap.

#### DNA sequencing of additional Cladophorales species and phylogenetic analysis

Sequence data were obtained from 9 additional Cladophorales species, representing the main lineages of the order (Supplementary Table 7). Total genomic DNA was extracted using a modified CTAB extraction protocol as described above, and sequenced using Illumina HiSeq 2000 technology (2×100 bp paired-end reads) on 1/5^th^ of a lane by Cold Spring Harbor Laboratory (Cold Spring Harbor, NY, USA). Quality of the reads from the sequenced libraries was assessed with FastQC 0.10.1. Low-quality reads (average Phred quality score below 20) were discarded and low-quality 3’ ends of the reads were trimmed with Fastx 0.0.13 toolkit. After trimming, reads shorter than 50 bp were discarded. Trimmed reads were assembled with CLC Genomics Workbench, MIRA and SPAdes 3.6.2^98^.

The taxonomic profiling of the contigs was performed with the following protocol: for each contig, sequence similarity searches were combined with the NCBI Taxonomy ID’s of the top ten BLAST hits in order to discriminate between eukaryotic and bacterial contigs and contigs with no similarity to known proteins ("NoHit"). Contigs classified as eukaryotic were further analysed to identify chloroplast contigs with a sequence similarity search using Tera-BLAST^TM^ (DeCypher, www.timelogic.com) against Chloroprotein_db. After chloroplast contig identification, the assembly that allowed the reconstruction of the highest number of full-length chloroplast genes was retained. An overview of the assembly metrics is reported in Supplementary Table 8, and the length distributions of the assembled contigs are reported in Supplementary Fig. 19.

Phylogenetic analysis was based on a concatenated alignment of 19 chloroplast protein-coding genes (*atpA*, *atpB*, *atpH*, *atpI*, *petA*, *petB*, *petD*, *psaA*, *psaB*, *psaC*, *psbA*, *psbB*, *psbC*, *psbD*, *psbE, psbF*, *psbJ*, *psbL*, and *rbcL*) from *Boodlea*, nine other Cladophorales species, 41 additional species of Archaeplastida, and 14 Cyanobacteria species (Supplementary Table 10). For each gene, DNA sequences were aligned separately using ClustalW codon alignment in Geneious. Gene alignments were concatenated and poorly aligned positions removed using the Gblocks server^99^, preserving the DNA codons, and using the least stringent settings. The resulting alignment was translated to obtain an amino acid alignment of 6,815 positions. Phylogenetic trees were inferred from the amino acid alignment using RAxML with cpREV + Γ4 model, using default parameters^100^. Branch support was assessed by bootstrapping with 500 replicates. Phylogenetic analysis was run on the CIPRES Science Gateway v3.3^101^.

## Data availability

DNA sequence data have been deposited to the NCBI Sequence Read Archive as BioProject PRJNA384503. The annotated chloroplast contigs of *Boodlea composita* and additional Cladophorales species were deposited to GenBank under accession numbers ***–***.

**Supplementary Information** is available in the online version of the paper.

## Acknowledgments

We thank Ellen Nisbett, Christopher Howe, Bram Verhelst, Sven Gould, Joe Zuccarello, and John W. La Claire II for help and advice. This work was supported by Ghent University BOF/01J04813, the Australian Research Council (DP150100705) to H.V., and the National Science Foundation (GRAToL 10136495) to J.L.B.

## Author Contributions

F.L., M.T., C.B., H.V., K.V. and O.D.C conceived the study. F.L. and C.B. provided chloroplast 454 data. A.D.C., K.V. and O.D.C provided the RNAseq and PacBio data. F.L. and J.M.L-B. provided genome data of additional Cladophorales species. A.D.C., F.L., K.A.B., J.J., K.V. and O.D.C analysed genomic data (including genomes assemblies, annotations, and phylogenetic analyses). A.D.C., F.L., K.A.B., M.T., J.J., H.V., K.V. and O.D.C wrote, and all authors edited and approved, the manuscript.

